# Scouter: Predicting Transcriptional Responses to Genetic Perturbations with LLM embeddings

**DOI:** 10.1101/2024.12.06.627290

**Authors:** Ouyang Zhu, Jun Li

## Abstract

This paper addresses the challenging problem of predicting transcriptional outcomes— the expression levels of all genes—in gene perturbation experiments and introduces a novel method called Scouter. By leveraging the capabilities of large language models and employing a neural network that facilitates easy training, Scouter overcomes key limitations of current approaches and accurately predicts the outcomes of single-gene and two-gene perturbations, reducing the error of state-of-the-art methods by half or more.

---

Gene perturbation experiments play a crucial role in biological and biomedical research due to their potential to reveal causal relationships and their direct medical and translational implications [1, 2]. As illustrated in Figure 1a, perturbing a gene—via knockout, enhancement, or suppression—typically alters the expression of many other genes due to intricate gene interactions and regulatory dynamics. These changes can be quantified using techniques such as Perturb-seq [3]. However, the high costs and practical constraints typically restrict the number of genes that can be individually perturbed in these experiments.

**Figure 1:**
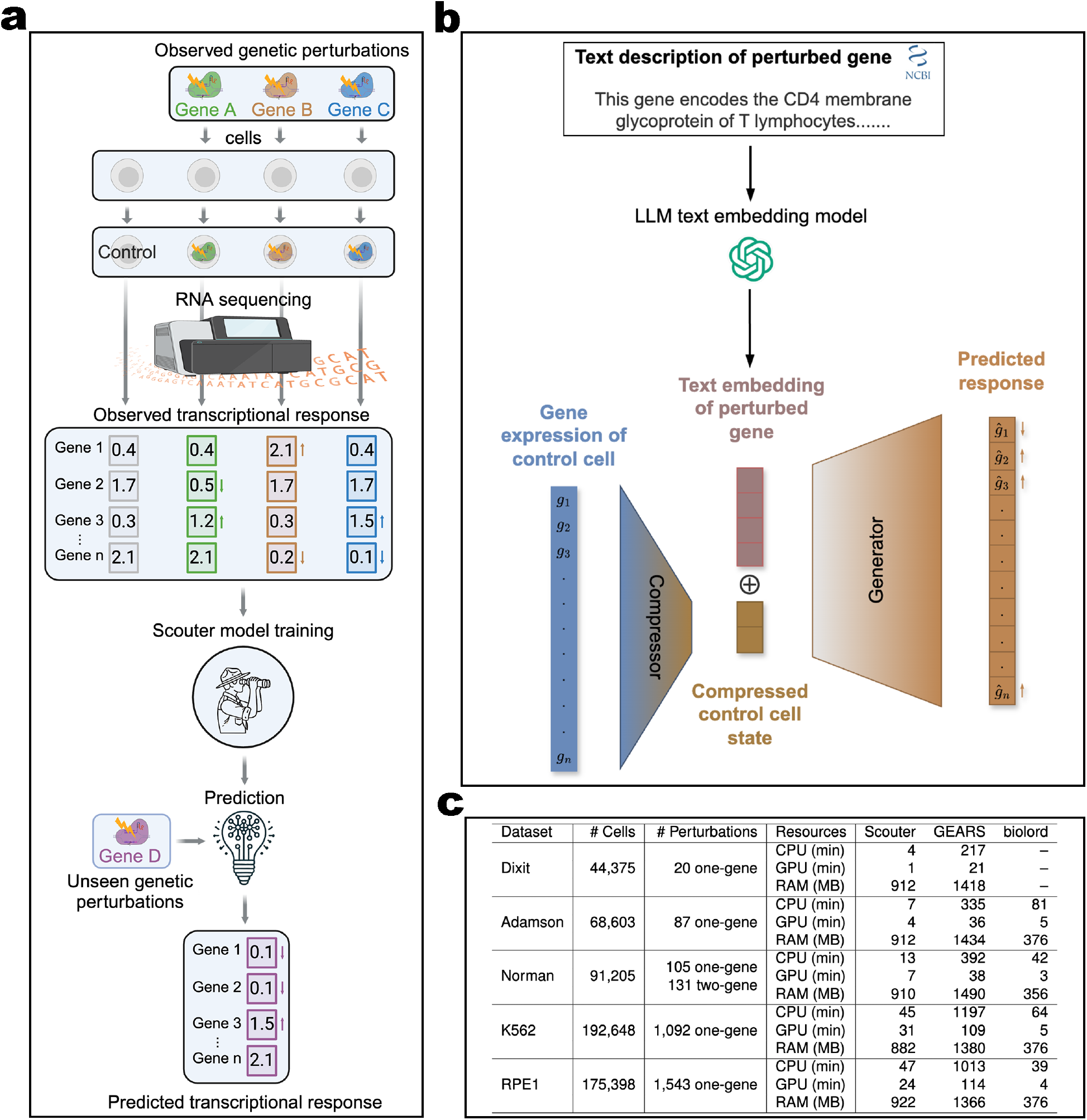
The problem Scouter addresses, the architecture of Scouter, and the datasets we use. **(a)** The perturbation experiments provide gene expression data for multiple single-gene perturbations. Methods such as Scouter learn from observed transcriptional responses and predict the responses of unseen perturbations. For clarity in this illustration, we focus on single-gene perturbations. Data from two-gene perturbations are also available and can be interpreted in a similar manner. Scouter is applicable to both single- and two-gene perturbations. **(b)** Scouter employs a neural network architecture that includes a compressor and a generator. **(c)** A summary of the datasets used and the computational time/resources required by different methods. Each dataset contains about 5,000 genes. Statistics for biolord on the Dixit data are not reported due to its exceptionally low predictive accuracy on this dataset.

The challenge of predicting the outcomes of unseen perturbation experiments (illustrated in Figure 1a) is significant. The primary issue is the high dimensionality of the desired prediction outcome—the expression levels of all genes—contrasted with the simplicity of the input: the identity of the perturbed gene, which is a single categorical variable. Although one could incorporate the expression data from an unperturbed (control) cell as input, this information remains constant across all experiments. The critical variable, the identity of the perturbed gene, remains singular and poses a significant challenge for encoding. Traditional methods such as one-hot encoding are inadequate because they cannot accommodate unseen perturbed genes without altering the input length, thereby rendering the model entirely non-functional for such predictions.

To address these challenges, any feasible algorithm must account for prior knowledge of gene-gene interactions and represent an unseen gene within the same space as seen genes. The currently available algorithms, GEARS [4] and biolord [5], both utilize gene regulatory networks informed by the Gene Ontology (GO) graph [6], where nodes represent genes and edges are weighted by the number of shared GO terms. These algorithms represent each perturbed gene by a combination of its nearest neighbors in the graph.

Despite these advancements, the reliance on the GO graph presents significant limitations. Firstly, the GO graph is sparse, meaning that only a small fraction of gene pairs share GO terms, which can compromise the accuracy and reliability of predictions. Secondly, the exclusion of some genes from the GO graph precludes predicting outcomes for perturbations involving these genes. Consequently, GEARS and biolord are unable to predict outcomes for all genes. Lastly, the use and extensibility of these algorithms are constrained by the specialized nature of machine learning techniques required for graph data.

In this paper, we propose a new method called Scouter, short for a tran**sc**riptional response predict**o**r for **u**nseen gene**t**ic p**er**turbtions with large-language-model (LLM) embeddings, that overcomes previous limitations and significantly enhances prediction accuracy. Scouter employs a fundamentally different approach to capture gene-gene interactions: the embeddings of genes generated by pre-trained LLMs. These embeddings, represented as long and dense numeric vectors, have been demonstrated in recent studies to capture complex regulatory relationships between genes more efficiently than traditional graphical models [7, 8, 9, 10, 11, 12]. Specifically, the gene embeddings used by Scouter, each of length 1,536, were sourced from the GenePT [11] paper. The generation of these embeddings did not require users to train their own models. Instead, they were produced at very low cost by utilizing OpenAI/ChatGPT’s “text-embedding-ada-002” model, which converts textual descriptions of genes into embeddings. These descriptions, such as the official NCBI gene descriptions, are rich in information about gene regulations, functions, and structures.

Scouter incorporates both the expression of all genes in the control cell and the ChatGPT embedding of the perturbed gene as input to a neural network that follows an compressor-generator framework, depicted in Figure 1b. Inspired by the concept of “disentanglement” stated in [13], Scouter first compresses the gene expression of the control cell into a short and dense vector. This vector is then concatenated with the text embedding of the perturbed gene, which subsequently serves as the input to the generator.

Scouter is evaluated on the five Perturb-seq datasets selected and pre-processed by the GEARS paper: Dixit [3], Adamson [14], Norman [15], Replogle K562 [16], and Replogle RPE1 [16]. The basic information of these datasets is provided in Figure 1c. These datasets each contain more than 40,000 cells, but the number of different perturbations can be as few as 20. This scenario makes training the model tricky: if one averages the expression profiles of all control cells to use as the input, and similarly averages the profiles of all cells with the same gene perturbation as the output, the training may suffer significantly due to an insufficient sample size. Therefore, unlike GEARS and biolord, Scouter randomly selects a control cell and a perturbed cell, using them as the network’s input and output for training, respectively. This approach, given *n*_0_ control cells in the data and *n*_*k*_ cells with gene *k* perturbed, yields a total number of training samples equal to *n*_0_ Σ_*k*_ *n*_*k*_, which is numerous. This strategy, combined with Scouter’s relatively simple network architecture, allows it to train effectively even on the challenging Dixit dataset.

Furthermore, despite the increased training sample size, Scouter maintains a reasonable memory footprint and demonstrates computational times comparable to those of biolord, both of which are more efficient than GEARS. Detailed metrics are presented in Figure 1c.

To quantitatively evaluate the results of different methods, we adopt the metrics used by GEARS and biolord (see Methods for detailed definitions): normalized mean squared error (MSE) and one minus normalized Pearson correlation coefficient (1-PCC) across the top 20 differentially expressed genes (DEGs). Both metrics favor small values. Figures 2a and 2b show the median (height of the bar) and the 50% confidence interval (ticks on the top of the bar) of these two metrics for each method across the five datasets. Clearly, Scouter achieves the lowest MSE and 1-PCC values in all five datasets, with substantial improvement over the other two methods: on average, Scouter’s MSE and 1-PCC are only about half of biolord’s (56.4% and 53.2%) and GEARS’s (49.6% and 48.0%).

**Figure 2:**
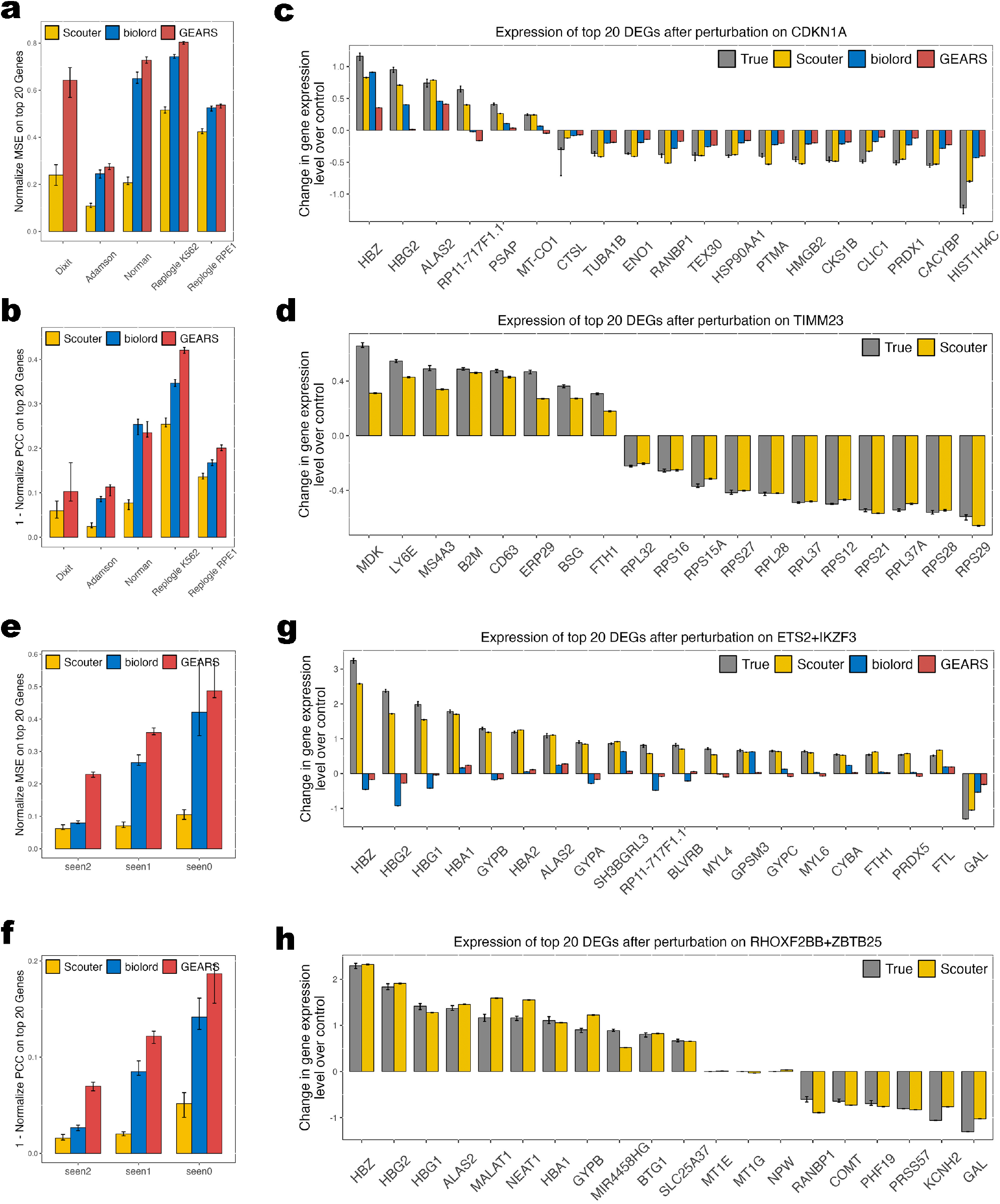
Predictions of different methods on real datasets. (Biolord’s results for the Dixit dataset are omitted due to exceptionally low predictive accuracy.) **(a-d)** One-gene perturbations. **(e-h)** Two-gene perturbations. **(a, e)** MSE on different datasets. **(b, f)** 1-PCC on different datasets. **(c, d, g, h)** Examples of true and predicted expressions for the top 20 DEGs.

Figure 2c showcases the true versus predicted expression profiles of the top 20 DEGs following perturbation of gene CDKN1A. Scouter’s predictions are more accurate than those of biolord and GEARS for 18 out of 20 genes (90%). Figure 2d focuses on the perturbation of gene TIMM23, which is absent from the GO graph, thus rendering biolord and GEARS unable to provide predictions. In contrast, Scouter continues to deliver precise predictions for the majority of these top DEGs.

Moreover, Scouter is capable of predicting transcriptional responses from the simultaneous perturbation of two genes. This functionality is implemented by summing the ChatGPT embeddings of the two genes and using the resultant vector as the input embedding. Figures 2e and 2f show the MSE and 1-PCC on the Norman dataset, which is the sole dataset among the five that includes two-gene perturbations. Following biolord and GEARS, the predictions are categorized based on the number of the two genes perturbed in the training data: seen2 (both were perturbed, but not simultaneously), seen1 (only one was perturbed), and seen0 (neither were perturbed). Scouter substantially outperforms biolord and GEARS, with its MSE and 1-PCC less than a half (43.4% and 39.8%) of biolord’s and less than a quarter (22.9% and 22.3%) of GEARS’s. Scouter’s performance is even more pronounced in two-gene perturbation cases compared to single-gene cases.

Figure 2g presents results for the dual perturbation of ETS2 and IKZF3, where Scouter accurately predicts the expression of all 20 genes. In contrast, both biolord and GEARS not only fail to predict the magnitude of the changes accurately but also make numerous errors in the direction of the changes. Figure 2h provides an example of a dual perturbation where the other models cannot make predictions due to limited GO graph coverage. Here, Scouter still offers accurate predictions for almost all genes.

In summary, Scouter markedly improves upon existing methods, reducing prediction errors by half for single-gene perturbations and by more than half for dual-gene perturbations. The efficacy of Scouter likely stems from the high-dimensional gene embeddings from LLMs, which are dense vectors packed with comprehensive gene information including but not limited to gene-gene interactions. Furthermore, Scouter’s uncomplicated network structure and its training strategy utilizing random selections of control and perturbed cell pairs have proven pivotal.

Moving forward, while Scouter currently extends single-gene perturbation predictions to two-gene scenarios using a straightforward summation of embeddings, exploring more sophisticated strategies will be a focus for future research. Additionally, the development of a zero-shot prediction method that does not rely on experimental perturbation data represents another significant direction.

## Methods

### LLM embeddings of gene text description

Predicting responses to unseen genetic perturbations represents a classical task of extrapolation in categorical variables, which is inherently challenging due to the lack of inherent order or distance among these variables. This makes it difficult for models to generalize from known to unknown genes. Motivated by the demonstrated effectiveness of LLMs in genomic research [12, 11], we propose utilizing embeddings of gene textual descriptions. By transforming the rich semantic content of textual descriptions into continuous vector representations with meaningful distances, we aim to capture semantic relationships between genes. This method allows the model to infer properties of unseen genes based on their descriptions, offering a scalable solution to the challenge of extrapolation.

The LLM embeddings considered in this work was provided in GenePT [11], which employs NCBI text descriptions of each gene and OpenAI’s “text-embedding-ada-002” model to generate the corresponding LLM embeddings [17] with a length of 1,536.

### Model architecture of Scouter

Scouter is a deep neural network model with a simple architecture that facilitates easy training. For a dataset containing *n* cells, with *n*_0_ control cells and *n* − *n*_0_ perturbation cells, each training epoch for Scouter processes *n* − *n*_0_ triplets: 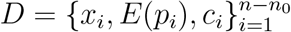. Here *x*_*i*_ is the gene expression vector of perturbed cell *i, p*_*i*_ represents the perturbed gene in cell *i, E*(*p*_*i*_) denotes the embedding vector of *p*_*i*_, and *c*_*i*_ is the gene expression vector of a randomly selected control cell.

Scouter incorporates a compressor *C* and a generator *D* (Fig. 1). Initially, *C* condenses the highly sparse vector *c*_*i*_ into a compact cell state *S*_*i*_ using a narrow bottleneck. Subsequently, *S*_*i*_ and *E*(*p*_*i*_) are concatenated and provided to *D*, which then generates the corresponding transcriptional response 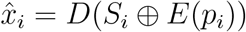.

To predict the response to a perturbation on gene *g*, Scouter takes as input the embedding *E*(*g*) and *K* random control cells 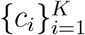 (with *K* set to 300), and returns 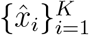. Then the average of 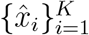 is output as the point estimator for the response prediction.

### Loss function of Scouter

Scouter is trained using a loss function known as the autofocus direction-aware loss, initially proposed by GEARS. This loss function comprises two components: the autofocus loss and the direction-aware loss. Consider a batch of *N* triplets 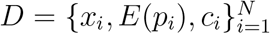, which includes *M* unique perturbations *t*_1_, *t*_2_, …, *t*_*M*_. The autofocus loss is defined as:

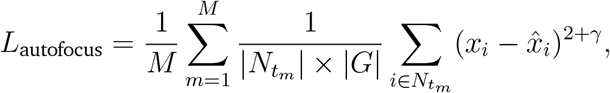

where |*N*_*tm*_| denotes the number of triplets with perturbation *t*_*m*_ and |*G*| is the length of the vectors *x*_*i*_ or 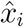. This loss penalizes discrepancies between the predicted response vector 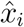 and the observed response vector *x*_*i*_. The parameter *γ* adjusts the focus of the penalty on larger discrepancies. On the other hand, the direction-aware loss is defined as:

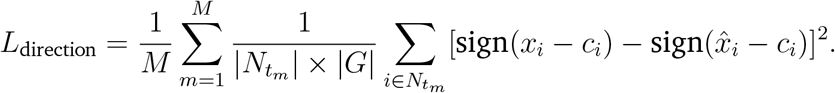

This component imposes additional penalties for errors in predicting the direction of changes. The total loss is then expressed as *L* = *L*_autofocus_ + λ*L*_direction_, where λ is the weighting factor for direction awareness.

### Data splits

Following the approach used in GEARS and biolord, we generated five different train-validation-test splits for the Adamson, Norman, Replogle K562, and Replogle RPE1 datasets, with 20% of the data set aside for testing, and the remaining 80% split into 90% for training and 10% for validation. Due to the small number of perturbations in the Dixit dataset, we generated 10 different splits using an 80:10:10 train-validation-test ratio. For each dataset, we used the validation set for early stopping and hyperparameter tuning, and evaluated the model’s performance as the average across all test sets.

### Benchmarks

We compared Scouter’s performance to that of biolord and GEARS. The evaluation was conducted using the settings provided in their respective reproducibility reposito-ries: https://github.com/nitzanlab/biolord_reproducibility/tree/main/scripts/biolord for biolord, and https://github.com/yhr91/gears_misc/blob/main/paper/fig2_train.py for GEARS.

### Metrics of prediction performance

Due to the fact that many genes do not exhibit significant differences between control and perturbed states, GEARS and biolord quantified model performance using normalized MSE and normalized PCC across the top 20 DEGs. Given the expression vectors of a perturbed cell and a control cell, denoted as *x*_*i*_ and *c*_*i*_, respectively, and the corresponding prediction 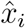:

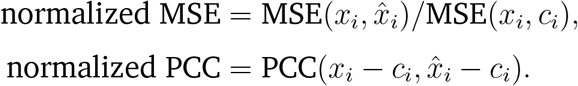

We adopted the same criteria, with the exception that we used 1 − normalized PCC instead of normalized PCC, so that both criteria favor smaller values.

### Selection of hyperparameters

For each dataset, we identified optimal hyperparameters by assessing the model’s average performance across the validation sets of all splits. Due to the large number of possible combinations, we fixed several hyperparameters: hidden layer sizes in the compressor at (2048, 512), compressor output layer size at 64, hidden layer size in the generator at 2048, dropout rate at 0, batch size at 256, learning rate decay factor at 0.9, maximum training epochs at 40, and patience at 5. We then conducted a grid search for the remaining hyperparameters within the following ranges: *γ* in the loss function at {0, 2}, λ in the loss function at {0.01, 0.05, 0.1, 0.5}, learning rate at {0.001, 0.005, 0.01}. The final hyperparameters selected were: *γ* at 0 for all datasets; λ at 0.5 for Replogle K562 and Replogle RPE1, 0.05 for Norman and Dixit, and 0.01 for Adamson; learning rate at 0.01 for Dixit, and 0.001 for all other datasets.

### Computing resources

We measured the maximum RAM usage of each method using the nvidia-smi command. For CPU runtime measurements, a MacBook Pro (2022) equipped with 16 GB of RAM and an Apple M2 chip (8-core CPU) was utilized. GPU runtime measurements were conducted on a server outfitted with a 32-core Intel(R) Xeon(R) Gold 6326 CPU, 256 GB of RAM, and an NVIDIA A40 GPU, which possesses 48 GB of GPU memory.

## Data availability

All datasets used in this study were obtained through the GEARS package, which provides preprocessed and thoroughly annotated data. The original datasets can be accessed via the Gene Expression Omnibus under the following accession numbers: Dixit: GSE90063; Adamson: GSE90546; Norman: GSE133344; Replogle K562 and Replogle RPE1: GSE146194. The GenePT gene embeddings are available at https://github.com/yiqunchen/GenePT.

## Code availability

Scouter is available as a Python package on GitHub at https://github.com/PancakeZoy/scouter.

## Notes

### Competing Interest Statement

The authors have declared no competing interest.

